# Polygenic adaptation from standing variation underlies rapid evolution under anthropogenic selection in an agricultural weed

**DOI:** 10.64898/2026.08.18.745463

**Authors:** Célia Neto, Augustin Baussay, Paul Neve

## Abstract

Herbicide resistance is among the clearest examples of rapid adaptation to intense anthropogenic selection. Yet, how the evolutionary origins and genetic architecture of resistance shapes its tempo and mode of evolution remain incompletely resolved. Here, we address these questions in *Alopecurus myosuroides* (blackgrass), Europe’s most widespread and economically damaging herbicide-resistant weed. We present the first genome-wide analysis of herbicide resistance in natural blackgrass populations, uniquely combining historical and contemporary populations collected before and after the onset of intensive herbicide use. This temporal framework provides novel empirical access to pre-selection genetic variation, enabling reconstruction of the tempo and mode of both target-site (TSR) and non-target-site resistance (NTSR) evolution across space and time. TSR mutations were not found in pre-herbicide populations and evolved recently through repeated, largely independent origins across Europe. NTSR, in contrast, has a polygenic architecture and is associated with a cluster of glutathione S-transferases (GSTs) with signatures of copy number variation, and broader stress-response genes. Most NTSR-associated alleles were already segregating in historical populations, consistent with rapid adaptation from standing genetic variation. Moreover, resistance-associated loci show signatures consistent with positive selection predating herbicide use, suggesting these stress and detoxification pathways were historically maintained by prior ecological selection and subsequently recruited under herbicide pressure. Together, these findings demonstrate that herbicide resistance encompasses contrasting genetic routes, with polygenic NTSR evolving largely through selection on standing variation, offering broader insights into the evolutionary dynamics of rapid polygenic adaptation under novel anthropogenic selection.

## Introduction

Understanding the evolutionary dynamics of adaptive traits is a central goal in evolutionary biology. Several fundamental questions determine the mode and tempo of rapid evolutionary responses to novel environmental stressors. Which genetic architectures underlie adaptation? Do adaptive traits arise from many, small-effect mutations or from a few large-effect variants? Is evolution parallel and/or convergent, and therefore predictable? Do adaptive traits arise from *de novo* or standing genetic variation? (Fisher, 1918; Provine, 2001; Barton and Keightley, 2002; Orr, 2005; Rockman, 2012; Lee et al., 2014; Barton et al., 2017; Barghi et al., 2020; Bomblies and Peichel, 2022).

Rapid evolution – especially under human-imposed selective pressures – provides a powerful system for studying how contemporary selection shapes evolutionary responses. Well-defined selective pressures and documented management histories enable explicit links between selection and observed evolutionary change (Hendry et al., 2017; Otto, 2018; Pelletier and Coltman, 2018). Classic examples include the evolution of industrial melanism in the peppered moth *Biston betularia* (Saccheri et al., 2008; Hof et al., 2011, 2016), drug resistance in pathogens (Palumbi, 2001; Laehnemann et al., 2014), and pesticide resistance in agricultural systems (Baucom, 2019; Hawkins et al., 2019; Gunn et al., 2024).

Agricultural systems impose recurrent, quantifiable and predictable selective pressures. As a result, pests – weeds, insects and pathogens – provide powerful model systems for studying rapid adaptation under strong anthropogenic selection with herbicide resistance in weeds being widely regarded as one of the clearest contemporary examples of “evolution in action” (Baucom, 2016; Kreiner et al., 2018). The repeated evolution on herbicide resistance across species (Palumbi, 2001; Baucom, 2019; Hawkins et al., 2019) has therefore become a powerful system for testing core evolutionary questions about genetic architecture, parallel adaptation, and the role of standing genetic variation (Délye et al., 2013a; Kreiner et al., 2018, 2019; Baucom, 2019; Van Etten et al., 2020; Comont et al., 2020; Kreiner et al., 2021, 2022b; Brunharo and Tranel, 2023; Cai et al., 2023; Kersten et al., 2023; Gupta et al., 2023). With increasing access to genomic resources for key weed species (Montgomery et al., 2024), it is now possible to unravel the evolutionary genomics of herbicide resistance across populations spanning a species range and contrasting herbicide selection histories.

Herbicide resistance evolves via two fundamentally distinct genetic architectures. Target site resistance (TSR) is typically monogenic and caused by specific point mutations in herbicide target genes. For two of the major herbicide target genes, acetolactate synthase (*ALS)* and acetyl co-A carboxylase (*ACCase)*, previous research has identified a well-defined, finite catalogue of single nucleotide polymorphisms (SNPs), each functionally and genetically validated, that result in herbicide resistance. These missense mutations have emerged repeatedly and independently across grass weed species worldwide (Heap, 2025; Powles and Yu, 2010). Non-target-site resistance (NTSR), on the other hand, is generally polygenic, involving multiple loci associated with metabolism, defence, and stress-response pathways across the genome. The precise genetic basis often varies among populations and species, and is therefore less repeatable and consequently less well characterized (Délye, 2013; Gaines et al., 2020). Because genetic architecture shapes the rate, repeatability, and predictability of adaptation (Barton and Keightley, 2002; Barghi et al., 2020), distinguishing the genomic bases of TSR and NTSR is central to understanding how resistance evolves (Kreiner et al., 2018; Baucom, 2019).

Experimental studies show that herbicide resistance can evolve within as few as three generations (Neve and Powles, 2005a; Powles and Yu, 2010), consistent with a major role for standing genetic variation. Theory predicts that adaptation from standing variation can proceed rapidly, since multiple pre-existing alleles are selected simultaneously, without the need to wait for *de novo* beneficial mutations (Hermisson and Pennings, 2005; Barrett and Schluter, 2008; Messer and Petrov, 2013; Boyle et al., 2017; Höllinger et al., 2019; Barghi et al., 2020). Characterizing these scenarios requires knowledge of pre-selection baselines and distribution of resistance-associated variants prior to widespread selection, which are rarely available in natural populations (Délye et al., 2010; Kreiner et al., 2021, 2022b; Kersten et al., 2023). Historical collections, including populations sampled before the onset of intensive herbicide use, are therefore uniquely valuable resources, offering a direct empirical window into pre-selection allele frequencies that no modelling or contemporary sampling approach can substitute.

In parts of Europe, herbicide resistance is frequent and widespread in *Alopecurus myosuroides* (blackgrass), a dominant grass weed in winter cereal systems (Moss, 2017; Hicks et al., 2018). TSR in blackgrass, particularly to ACCase inhibitors, has evolved repeatedly through 14 independent missense mutations at the herbicide target gene. These substitutions are now firmly established, functionally and genetically validated in prior work, and create a largely monogenic architecture, with strong molecular parallelism, documented repeatedly across European populations (Menchari et al., 2006, 2007; Délye et al., 2010; Kersten et al., 2023). In comparison, NTSR remains poorly characterized at the population level, since research has largely relied on laboratory-derived intercross populations or small samples of wild individuals, limiting broader inference about its genetic basis in natural populations (Délye et al., 2010; Délye, 2013; Dixon et al., 2021; Cai et al., 2023).

Here, we investigate the genetic architecture and evolutionary dynamics of herbicide resistance in blackgrass across space and time, leveraging a unique collection of contemporary and historical populations, spanning from the Middle East (blackgrass’ putative native range) and throughout Europe. We integrate genome-wide mapping, demographic analyses and selection scans to (i) resolve the origin and evolutionary history of TSR; (ii) characterize the genetic architecture of NTSR; and (iii) assess the relative contributions of standing vs *de novo* variation in the evolution of TSR and NTSR. Bridging evolutionary theory, and population and quantitative genomics, we infer the origin, repeatability, architecture and spatiotemporal dynamics of herbicide resistance evolution in blackgrass.

## Material and Methods

All detailed scripts are available at https://github.com/celianeto/blackgrass_HR. All statistical analyses and plotting were conducted in R v4.4.1 (R Core Team, 2021).

## Population collection and phenotyping

We assembled a collection of 173 blackgrass (*Alopecurus myosuroides*) populations from across the species’ distribution in the Middle East (putative native region) and Europe (Supplementary Table 1). Of these, 163 were collected directly from agricultural fields between 2013 and 2021, while ten populations were obtained from the USDA-ARS Germplasm Resources Information Network (GRIN, https://www.ars-grin.gov) seed stock. Here, “population” refers to a bulk seed lot originating from multiple plants at the same location. For contemporary populations, seeds were harvested by diverse collaborators and colleagues for the purposes of characterizing field-scale herbicide resistance in agricultural fields. Each population is composed by seeds from multiple plants within each field and bulked. Seeds used in this study were randomly drawn from this bulk sample. Historical samples obtained from the USDA-ARS GRIN collection derive from a single propagation of field-collected material. Original seeds collected from multiple plants in the field were grown and allowed to freely intercross (as *Alopecurus myosuroides* is an obligate outcrosser), as they would under natural conditions. Then, seeds from 100 distinct plants were harvested, combined, and maintained as bulk seed lots. Seeds provided by the USDA-ARS GRIN were from these bulk seed lots. The ten USDA-ARS GRIN populations consist of eight populations collected between 1953 and 1977 and two populations collected in 1997. Throughout the paper, *historical* refers to populations collected before 1977.

Blackgrass seeds from the 173 populations were incubated for one week at 17°C day / 11°C night with a 12h photoperiod in Petri dishes containing 4 mL of 2 g/L KNO_3_. Ten germinated seedlings per population were then transplanted into plant pots containing a standard soil potting mixture and maintained for two weeks under greenhouse conditions (22°C day / 18°C night) with regular watering. At the 2-3 leaves stage, all individuals were sprayed with a commercial formulation of the ACCase-inhibiting herbicide fenoxaprop-p-ethyl. The herbicide was applied at a standard field rate (69 g ai ha^-1^), using an experimental track sprayer, delivering a water volume of 133 Lha^-1^ at a pressure of 3 bar. Plant survival was scored two weeks after spraying, with individuals classified as resistant (alive with active growth post herbicide application) or susceptible (dead with no plant growth post herbicide application). Resistance at the population level was quantified as the proportion of surviving plants.

### Genomic data

Sixty-four representative populations (four Middle Eastern historical, four European historical, seven Middle Eastern contemporary, and 48 European contemporary) were selected for whole-genome pooled sequencing (Supplementary Table 1). For each population, 25 individuals were grown in a glasshouse and sampled for pool-sequencing. For each of those 25 individuals, 2 cm of a young leaf were cut and pooled. DNA was then extracted per pool, i.e., per population using the DNeasy Plant Mini Kit (Qiagen) following the manufacturer’s protocol. Genotyped individuals were sampled independently of those used for phenotyping but from the same bulk seed collections, providing independent population-level phenotypes and genotypes. Paired– end libraries (2×150 bp) were prepared and sequenced on an Illumina NovaSeq platform at Novogene, UK. Each population pool was prepared as a separate sequencing library and processed independently throughout read mapping, variant calling, and allele frequency estimation.

Raw reads were quality-trimmed using Trimmomatic (– “LEADING:5” – “TRAILING:5” – “SLIDINGWINDOW:4:20” – “MINLEN:50”) (Bolger et al., 2014), and then aligned to the *A. myosuroides* reference genome (NCBI version: JAPCYS010000000) (Cai et al., 2023) using the maximal exact matches algorithm in bwa (*bwa mem*) (Li, 2013), and default parameters. Alignment files were processed further with samtools (Danecek et al., 2021): reads were filtered for quality (*samtools view –q 20*) and sorted (*samtools sort*). Duplicate reads were marked and removed using picard (http://broadinstitute.github.io/picard). Per population bam files were combined into a single mpileup file using *samtools mpileup*.

Allele count data were generated using PoPoolation2 (Kofler et al., 2011), converting mpileup files to sync format, using the script mpileup2sync.jar. Sync files were further converted to BayPass-compatible input using the functions *popsync2pooldata()* and *pooldata2genobaypass()* from the poolfstat R package (Hivert et al., 2018), filtering for SNPs only, minimum read count of five per population (argument *min.rc = 5*) and minimum minor allele frequency of 0.05 (argument *min.maf = 0.05*). The output files were then used as input for downstream analyses in Treemix (Pickrell and Pritchard, 2012) and BayPass (Gautier, 2015). At the end, 103 888 478 SNPs were kept.

### Population characterization

The populations used here originated from multiple sources (USDA and collaborators), and the exact number of individual plants contributing to each seed bulk was therefore unknown. While contemporary populations were known to be derived from multiple individuals, equivalent information was not available for historical populations. Because this uncertainty could influence downstream analyses and interpretations, we first genetically characterized all populations to assess whether they represented large, well-admixed samples capturing existing phenotypic and genetic diversity, or whether they might instead consist of a limited number of related individuals. To this end, we quantified genetic diversity within populations using Watterson’s θ, as implemented in *npstat* (Ferretti et al., 2013), and tested for differences between historical and contemporary populations across regions using a linear model (*lm()* function in R).

### *ACCase* genotyping

Acetyl-coenzyme A carboxylase (*ACCase*) is the target enzyme for ACCase-inhibiting herbicides, including fenoxaprop-p-ethyl (Délye, 2005). Fourteen well-characterized resistance-conferring missense mutations have been reported across different grass species (Table 1) (Powles and Yu, 2010; Beckie and Tardif, 2012; Kersten et al., 2023). Because causality for these variants is established by prior work, we focused our TSR analysis on screening for this defined set of mutations. Allele frequencies (AF) for each mutation in each of the 64 sequenced populations were calculated as the proportion of reads containing the resistance allele relative to total coverage at each site, from the sync file, using the package poolfstat (Hivert et al., 2018) in R.

**Table 1.** *ACCase* mutations in European blackgrass. Columns show, from left to right, the previously described amino acid changes in *ACCase*, the number of populations they were found in in this study, the average frequency of each mutation across populations, the frequency range in all populations, and the p-value for the clustering test.

| Mutation | Number of populations | Average frequency in populations | Frequency range | Clustering (p-value) |
| --- | --- | --- | --- | --- |
| Ile1781Leu | 17 | 0.36 | 0.03-0.83 | 0.83 |
| Ile2041Asn | 7 | 0.19 | 0.03-0.55 | 0.56 |
| Asp2078Gly | 6 | 0.22 | 0.03-0.5 | 0.84 |
| Trp2027Cys | 5 | 0.16 | 0.05-0.34 | 0.03 |
| Ile1781Val | 3 | 0.42 | 0.12-0.93 | 0.93 |
| Gly2096Ala | 3 | 0.10 | 0.03-0.18 | 0.32 |
| Ile1781Thr | 2 | 0.06 | 0.04-0.07 | 0.61 |
| Trp1999Leu | 1 | 0.24 | - | - |
| Ile2041Val | 0 | - | - | - |
| Cys2088Arg | 0 | - | - | - |
| Trp1999Cys | 0 | - | - | - |
| Gly2096Ser | 0 | - | - | - |
| Ile1781Ala | 0 | - | - | - |
| Trp1999Ser | 0 | - | - | - |

### *ACCase* mutations’ origin and spread

We reconstructed population relationships at the *ACCase* locus (± 5 kb) using Treemix v1.13 (Pickrell and Pritchard, 2012). The input file was created using the functions *popsync2pooldata()* and *pooldata2genobaypass()* in the R package poolfstat (Hivert et al., 2018) on the genobaypass file, as described above. Treemix was run with 1000 SNPs grouped per window to account for linkage disequilibrium (flag *-k 1000*) (Pickrell and Pritchard, 2012) and trees were visualized in R using the ape package (Paradis and Schliep, 2019).

We then tested whether each resistance mutation exhibited statistically significant clustering on the tree, indicating a potential single origin. Phylogenetic clustering was assessed using a permutation-based mean pairwise distance (MPD) approach (Webb et al., 2002; Kembel et al., 2010). MPD was calculated as the average pairwise patristic distance among all populations carrying a given mutation, using the *cophenetic.phylo()* function from the ape package (Paradis and Schliep, 2019) in R. Statistical significance was determined by comparing the observed MPD to a null distribution generated from 10 000 random permutations in which the mutation was randomly assigned to the same number of tips on the tree. A one-sided p-value was computed as the proportion of permuted MPD values less than or equal to the observed value.

### Mapping the genetic basis of non-target-site resistance

We identified loci associated with non-target-site resistance (NTSR) using a genome-wide association (GWAS) analysis in BayPass (Gautier, 2015), applied to whole-genome pool-seq data from 48 contemporary European populations (Supplementary Table 1). The proportion of surviving plants per population following herbicide treatment was used as a quantitative proxy for resistance and served as the phenotype in the analysis. We implemented the standard covariate model (flag *-efile*). To explore genetic relationships among the 48 European populations, we performed principal component analysis (PCA) on the scaled covariance matrix of population allele frequencies, estimated by BayPass. PCA was visualized using the *plot.omega()* function in R from BayPass. The matrix of population allele frequencies captures the shared demographic history amongst populations and can be used for inferring population structure (Gautier, 2015).

To account for linkage disequilibrium, we applied the local score approach to the uniformly distribution of p-values and calculated Lindley scores (Bonhomme et al., 2019) from the BayPass output. Significant intervals were defined as regions exceeding the local score threshold, and genes located 10 kb upstream or downstream of these regions were considered candidate genes.

In addition to association statistics, BayPass provides the effect size (beta) for each variant, its allele frequency per population (corrected for population history), and the XtX statistic (see below). The allele frequency data were used to evaluate the geographical distribution of resistance-associated variants.

### Signals of selection

To test whether resistance-associated loci exhibited signatures of selection, we selected the variant with the largest estimated effect size within each significant region as the representative of that region. We then compared the distribution of population genetic statistics for this representative set of variants against the genomic background. Specifically, we calculated (i) Watterson’s θ, as a measure of diversity, using npstat (Ferretti et al., 2013), in 1 Mb windows (flag *-l 1000000*); and (ii) XtX, as provided directly by BayPass (Gautier, 2015). Statistical significance was assessed by comparing the distributions of resistance-associated loci against the genomic background using Kolmogorov–Smirnov tests (R function *ks.test()*).

XtX serves as a measure of adaptive differentiation and can be interpreted as a variant-specific F_ST_ explicitly corrected for the scaled covariance of population allele frequencies (Gautier, 2015). This analysis was conducted on the 48 European contemporary set and repeated in the set of historical populations alone to determine whether selection signals predated intensive herbicide use.

### Robustness analyses of allele frequency change

To confirm that observed allele frequency shifts at candidate NTSR loci reflect genuine selection rather than sampling artefacts, two robustness analyses were performed. First, contemporary resistant populations were downsampled to match the number of historical susceptible populations (n = 8) across 100 resampling replicates, and allele frequency changes at candidate loci were recalculated across the susceptibility-resistance continuum. No effect of sampling number was detected (Supplementary Figure 9). Second, 100 sets of 76 genome-wide variants with initial allele frequencies matching those of the candidate loci were randomly sampled, generating a null distribution of expected drift-driven shifts. Observed candidate loci shifts were then compared to this null distribution using a Kolmogorov–Smirnov test (*ks.test()*).

### Prediction of herbicide resistance

We also tested whether the set of resistance-associated variants could predict resistance phenotypes across populations. We modelled the survival proportion of each of the 48 European contemporary populations as a function of allele frequencies at the candidate variants identified. Prediction was performed using a Bayesian generalized linear model implemented with the *train()* function in the caret package (Kuhn, 2007), with 10 populations randomly selected as the training set and the remaining populations used for validation. This procedure was repeated 100 times, each with different training and validation sets. Predictions for validation populations were generated using the *predict()* function, and model performance was evaluated as the Spearman correlation between observed survival and the median predicted values across replicated runs. Missing genotype values were treated as absence of the allele (set to zero).

### Glutathione S-transferases as candidates

To investigate the region of interest on chromosome 3, we used the GenomeBrowser tool provided by the International Weed Genomics Consortium via WeedPedia (https://weedpedia.weedgenomics.org), where the blackgrass reference genome from Cai et al. (2023) is hosted. Sequence alignment and similarity among the five glutathione S-transferases (GSTs) genes were assessed using BLAST (NCBI; https://blast.ncbi.nlm.nih.gov/Blast.cgi), with similarity estimated using the Fast Minimum Evolution method.

Coverage across the target locus was quantified from the sync file (see above) using the poolfstat package (Hivert et al., 2018) in R, in 1 kb windows, and values were normalized per pool relative to mean coverage across chromosome 3.

## Results

### Herbicide resistance in blackgrass evolved recently in Europe

Across all populations, genetic diversity (Watterson’s θ) varied between 0.015 and 0.039. To account for heterogenous spatiotemporal sampling protocols, we modelled genetic diversity as a function of collection period, geographic origin, and their interaction (linear model, adjusted R^2^ = 0.61). While geographic origin (mean _Middle East_ = 0.028, mean _Europe_ = 0. 017) had a significant effect on diversity (p-value = 3.77×10^-5^), neither collection period (mean _historical_ = 0. 021, mean _contemporary_ = 0.022) nor the interaction between the two terms was significant (p-value = 0.524 and 0.619, respectively), indicating that historical and contemporary populations from the same region contain comparable levels of genome-wide diversity. This suggests that historical populations from the USDA-GRIN germplasm collection maintained genetic diversity comparable to contemporary field collections, providing a suitable representation of existing genetic variation prior to herbicide selection.

Across the sampled distribution, resistance to acetyl-CoA carboxylase (ACCase) inhibiting herbicides, measured as survival to a field relevant dose of fenoxaprop-p-ethyl (a widely used post-emergence herbicide for the control of annual and perennial grasses), showed a clear geographic pattern (Figure 1A). Of the 173 geographically diverse blackgrass populations tested, 16% (n = 26) were susceptible, with no individuals surviving herbicide exposure, while 56% of the populations showed survival rates (resistance) above 80% (Supplementary Figure 1). Populations from central and northern Europe were typically highly resistant, whereas populations from the Middle East – where blackgrass is likely native – were largely susceptible (mean _Middle East_ = 0.07, mean _Europe_ = 0.72, Wilcoxon test, p-value = 7.24×10^-9^). Stratifying populations by time of collection revealed that historical populations, collected before the intensification of herbicide use in the 1970s, were far more susceptible to ACCase herbicides than contemporary populations (mean _historical_ = 0.03, mean _contemporary_ = 0.69, Wilcoxon test, p-value = 4.58×10^-5^; Figure 1B). Only two individuals, one collected in Turkey in 1953 and another in Spain in 1963, out of 80 historical plants screened, survived herbicide application. Although unexpected, similar low frequency, pre-selection resistance to ACCase herbicides has been reported previously: up to 2.6% of individuals in wild *Lolium rigidum* populations from Australia, which had not been exposed to herbicides, exhibited resistance (Neve and Powles, 2005b).

**Figure 1.**
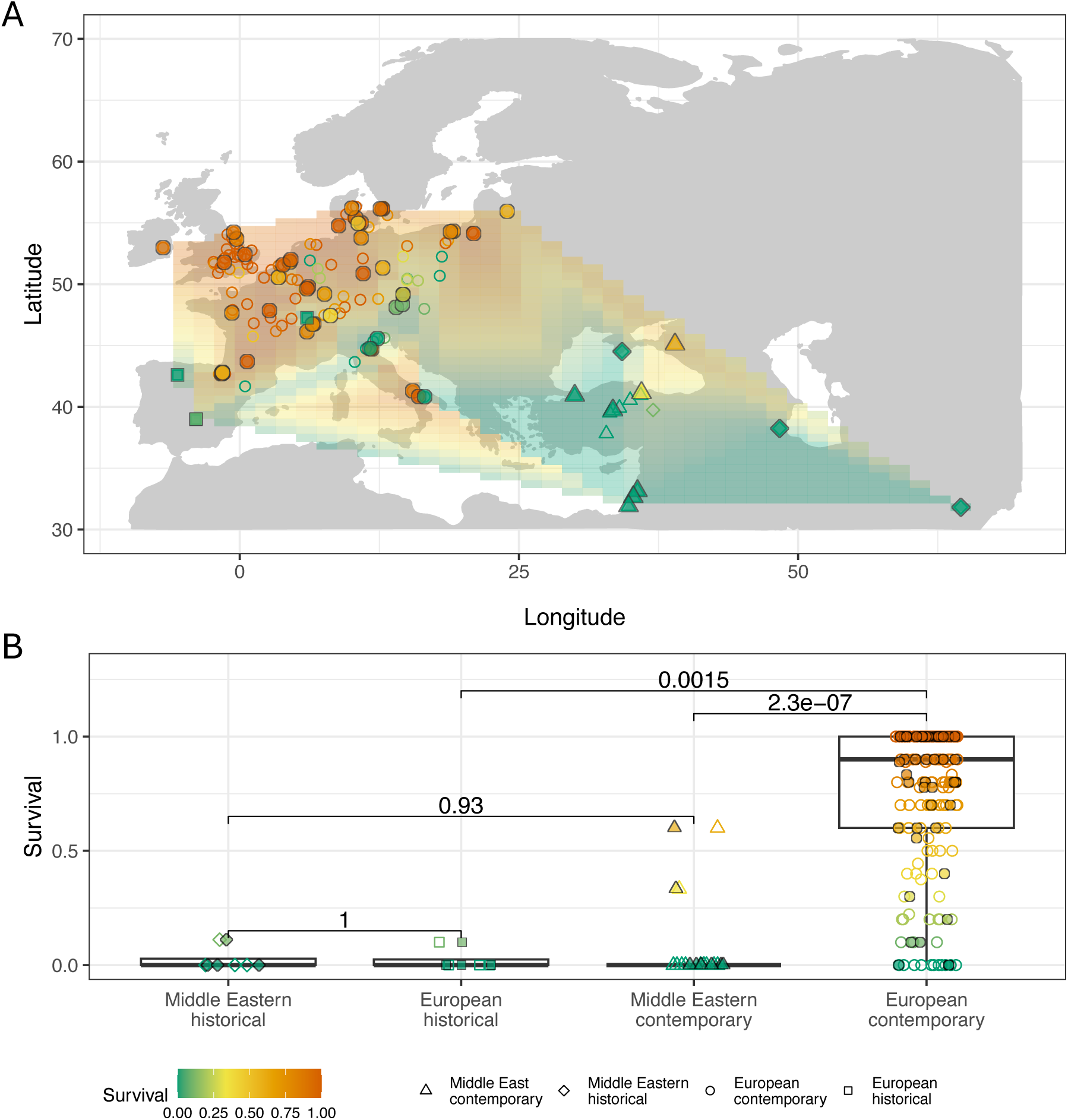
Recent evolution of herbicide resistance in blackgrass in Europe. A. Geographical distribution of populations used in this study, coloured according to survival to fenoxaprop per population. The shaded background represents the interpolated geographical distribution of the trait, with colours following the legend. Shapes reflect place and time of collection, according to the legend. B. Boxplot showing phenotypic distribution (survival to fenoxaprop, y-axis) per population category (x-axis). Categories reflect region of origin and time of collection (*historical* refers to populations collected before the Green Revolution, in the 1970s). Each point represents one population, with colour representing survival and shape population categories, as shown in the legend. Throughout the figure, filled dots mark sequenced populations. P-values result from the Wilcoxon text.

**Supplementary Figure 1.**
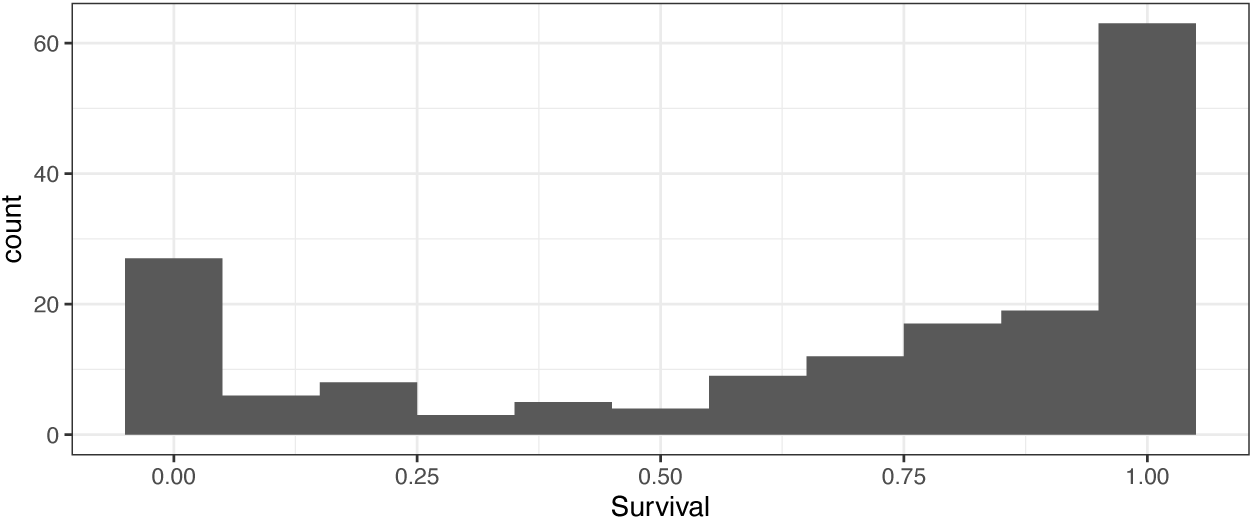
Phenotypic distribution for the 173 blackgrass populations screened. X-axis shows proportion of survival per population (10 individuals screened).

Overall, European contemporary populations – mainly from northern Europe – were significantly more resistant than both historical Middle Eastern populations (mean _historical Middle East_ = 0.03, Wilcoxon test, p-value = 0.002) and historical European populations (mean _historical Europe_ = 0.03, Wilcoxon test, p-value = 0.002; Figure 1B). These results indicate that herbicide resistance has evolved recently within the non-native range of blackgrass in central and northern Europe.

### ACCase resistance mutations have recently expanded across Europe

Next, we investigated the genetic basis of herbicide resistance in blackgrass. We sequenced 64 representative populations (Supplementary Table 1) and first examined target-site resistance (TSR) by screening for known resistance mutations in *ACCase* – the target gene of ACCase-inhibiting herbicides. We genotyped the fourteen previously validated functional *ACCase* mutations found across multiple grass weed species, including blackgrass (Table 1) (Vila-Aiub et al., 2009; Gaines et al., 2020; Kersten et al., 2023).

Across our 64 sequenced populations, we found eight of the 14 previously described *ACCase* mutations (Table 1). Ile1781Leu was the most frequent, occurring in 17 populations with allele frequencies ranging between 0.03 and 0.83. Trp1999Leu was the rarest, present in only one population from Luxembourg at 0.24 frequency. None of the mutations identified were fixed in any population, and their frequencies ranged from 0.03 to 0.93, with an overall average frequency of *ACCase* target-site mutations of 0.26.

Although most mutations were widespread, some *ACCase* mutations showed clear geographical structuring, with the greatest diversity of distinct mutations observed in northern France and the Benelux region (Figure 2A, Supplementary Figure 2B). Specifically, Trp2027Cys was found in northwestern Europe, Gly2096Ala spanned western Europe from the Pyrenees to Denmark, Ile1781Val occurred only in southern Italy and the Pyrenees, and Ile1781Thr was restricted to the Netherlands/northwest Germany. These patterns may reflect independent origins of the same mutation and subsequent dispersal, and/or fitness trade-offs that influence the relative costs and benefits of mutations in specific environments.

**Figure 2.**
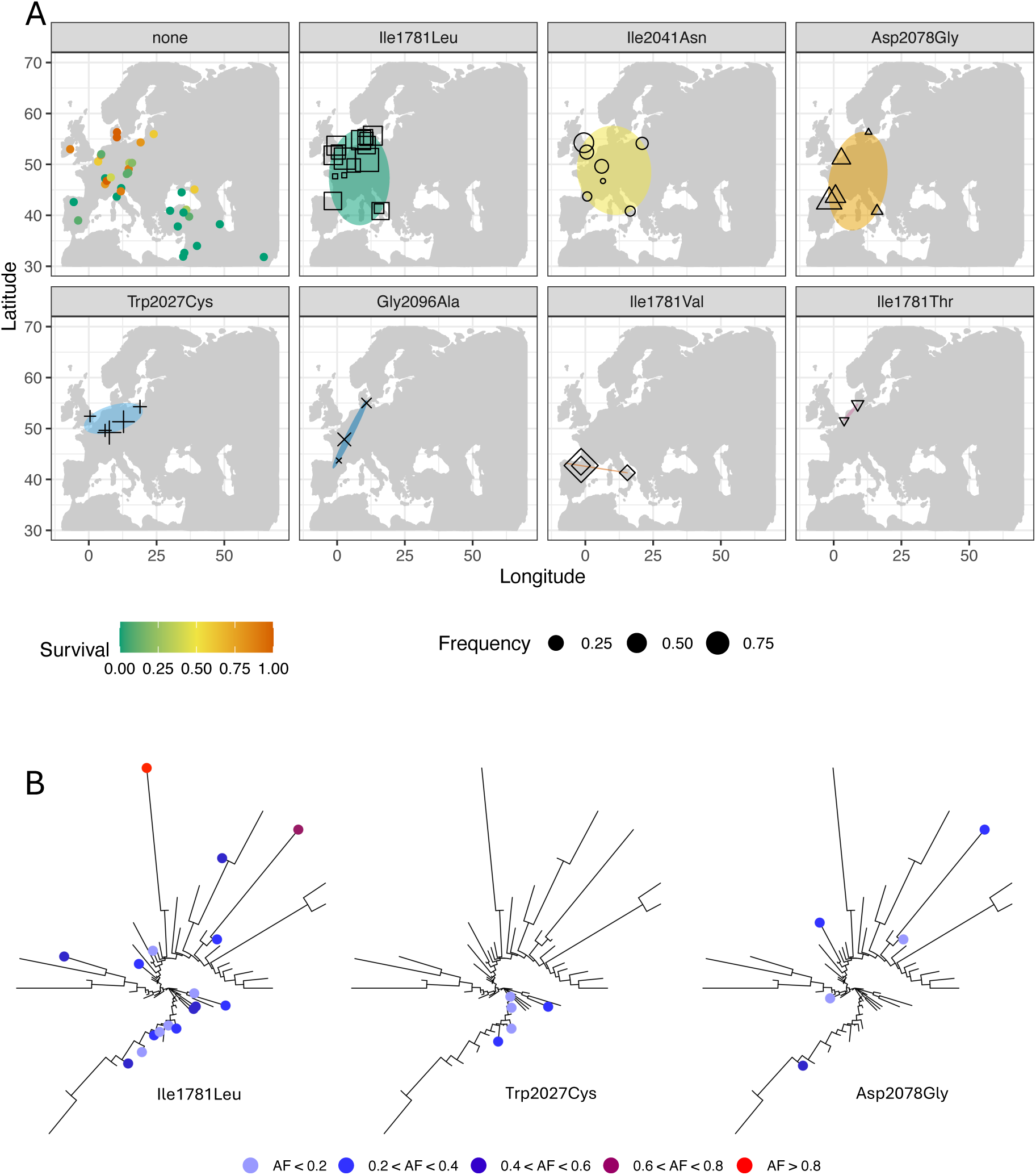
Origin of *ACCase* mutations in blackgrass. A. Geographical distribution of the different mutations found in *ACCase* in this study in the 64 sequenced populations, across the blackgrass distribution. Ellipses mark the distribution of the corresponding mutation, labelled on top of each facet. Symbol size corresponds to the frequency of that mutation in each population. Each mutation is represented by a different symbol for clarity. *None* refers to populations with no detectable *ACCase* mutations. These populations are coloured by survival, following the legend. B. *ACCase* locus maximum-likelihood trees for the 64 populations sequenced. In each tree, different mutations are marked, as shown by the labels. Each tip with a dot marks a population where the corresponding mutation was detected. The dot is coloured according to the frequency of that mutation in that population, following the legend (AF = allele frequency). A tip with no dot represents a population without that *ACCase* mutation.

**Supplementary Figure 2.**
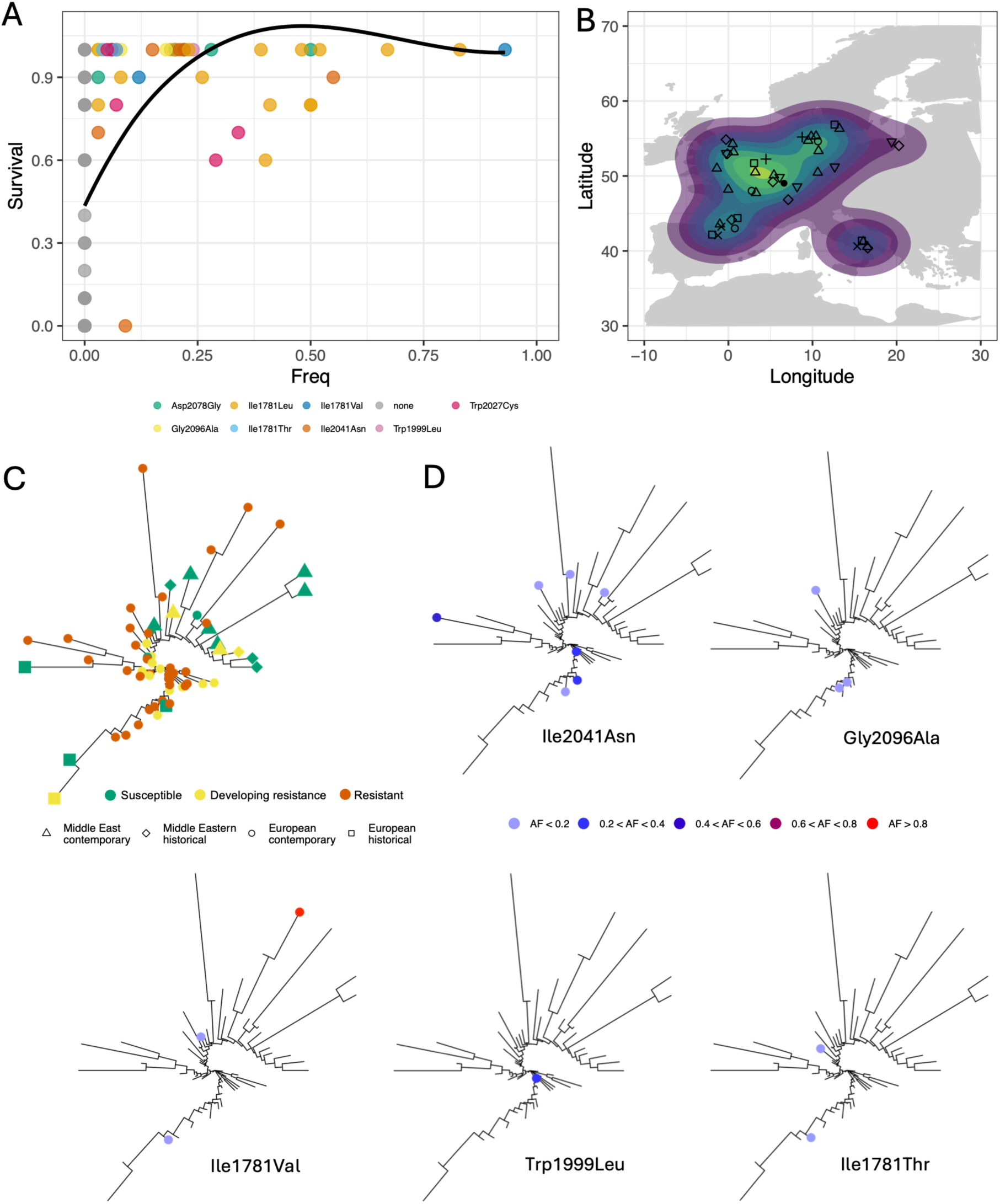
Origin of *ACCase* mutations. A. Correlation between allele frequency of each of the *ACCase* mutations found (x-axis) and survival (y-axis) per population (each dot). Mutations are coloured as shown in the legend. The black trend line comes from a Loess fit. B. Map of Europe showing the density of distinct *ACCase* mutations: lighter colours represent geographical regions with higher number of different mutations. Dots represent populations carrying specific mutations as in Figure 2A. C. *ACCase* tree of the 64 populations. Each tip is coloured by survival (susceptible: survival = 0; resistant: survival > 0.8; developing resistance: anything in between) and shapes show the different population categories. D. *ACCase* locus maximum likelihood tree. In each tree, a specific *ACCase* mutation is marked, as shown by the labels, by a dot coloured based on the allele frequency of that mutation in that population.

The six previously described *ACCase* mutations we did not find (Ile2041Val, Cys2088Arg, Trp1999Cys, Gly2096Ser, Ile1781Ala, and Trp1999Ser) were also not detected in a previous study with other European blackgrass populations (Kersten et al., 2023). Compared to that study, we found an additional resistance-endowing mutation, Gly2096Ala, in three populations (two in France, one in Denmark), at low frequencies.

No known *ACCase* mutations were identified in thirty-five of the sequenced populations (55%), despite some of those populations exhibiting phenotypic resistance to fenoxaprop (mean survival across all populations with no *ACCase* mutations = 0.33; Figure 2A). More importantly, no *ACCase* mutation was detected in any of the Middle Eastern or historical populations (Figure 2A), suggesting these mutations rose in frequency, in the last ∼50 years, in Europe.

### Multiple origins and gene flow underlie ACCase resistance in Europe

To assess how population history and dispersal might have shaped the distribution of *ACCase* mutations, we analysed their putative origin and spread using a maximum likelihood tree at this locus (± 5 kb) and mapped each mutation onto it (Figure 2B, Supplementary Figure 2C and 2D). Populations sharing mutations with a single origin are expected to be more clustered on the tree than by chance (p-value < 0.05; Table 1). While most *ACCase* mutations were found in populations distributed across the tree, suggesting multiple origins in agreement with their geographical spread, the clustering of some mutations was more discrete. For example, the most common mutation, Ile1781Leu, was widely distributed across the tree (clustering test, p-value = 0.83), potentially indicating multiple origins. Within the branch where it occurs most frequently, however, we observed significant clustering (clustering test, p-value = 0.03), suggesting a shared history for that mutation in those particular populations. In contrast, Trp2027Cys was mainly confined to a single branch (clustering test, p-value = 0.03), this being consistent with a single origin, followed by dispersal among populations via migration and/or gene flow. These results suggest that different ACCase*-*resistance-conferring mutations likely arose through (i) multiple independent origins (e.g., Asp2078Gly), (ii) a single origin followed by migration and/or gene flow (e.g., Trp2027Cys), or (iii) a combination of both processes (e.g., Ile1781Leu).

Overall, *ACCase* mutations explained 37% of the phenotypic variation observed in herbicide resistance (linear model, R^2^ = 0.37, p-value = 5.95×10^-9^; Supplementary Figure 2A), with 21 populations showing some level of resistance but no *ACCase* mutations. These results indicate that high resistance levels can occur in the absence of known TSR variants and suggest a major role for non-target-site resistance (NTSR) (Délye et al., 2011; Délye, 2013; Comont et al., 2020; Gaines et al., 2020; Kreiner et al., 2021; Kersten et al., 2023).

### Polygenic NTSR evolved via standing genetic variation

To explore the genetic architecture of putative NTSR, we performed a genome-wide association analysis (GWAS) on 48 contemporary European populations. By restricting the analysis to this subset, we minimized confounding effects from deeper population structure, private alleles, and historical demographic differences (Supplementary Figure 3). After correcting for linkage disequilibrium using the local score approach (Bonhomme et al., 2019), we identified 76 significant genomic regions associated with herbicide resistance (Figure 4A, Supplementary Table 2). For each QTL, the SNP with the largest estimated effect size was selected as the representative variant and polarized relative to the reference genome, which was derived from a susceptible individual (Cai et al., 2023), so that all inferences refer to the alternative allele. A predictive Bayesian generalized linear model demonstrated that these 76 representative SNPs were sufficient to predict herbicide resistance in contemporary European populations with high accuracy (Spearman’s ρ = 0.90, p-value < 2.2×10^-16^; Supplementary Figure 4), indicating that these variants capture a substantial proportion of the genetic signal underlying variation in NTSR in Europe.

To further characterize the genetic architecture of NTSR, we examined effect sizes and allele frequencies of the representative SNPs. These loci were characterized by large effects (mean |β| = 0.095), translating to an average 32% increase in herbicide survival per allele (in the model, maximum β is 0.3), indicating a polygenic architecture with large effect variants distributed across the genome. Median allele frequency across populations and effect size per locus were not correlated (Spearman’s ρ = –0.07, p-value = 0.56; Supplementary Figures 5 and 6), suggesting that both common and population-specific variants contribute to resistance. The spatial distribution of alleles reflected this pattern (Figure 3A, Supplementary Figures 6 and 7). Some alleles were at intermediate frequencies across Europe but reached very high or very low frequencies locally (Figure 3A, a-b). Others were consistently very frequent (Figure 3A, c-d) or consistently rare (Figure 3A, e-f) with sporadic local deviations. These patterns imply that NTSR arises from population-specific combinations of alleles at varying frequencies, producing a continuous, cumulative, dimmer-like response – in contrast to the discrete switch-like behaviour of TSR.

**Figure 3.**
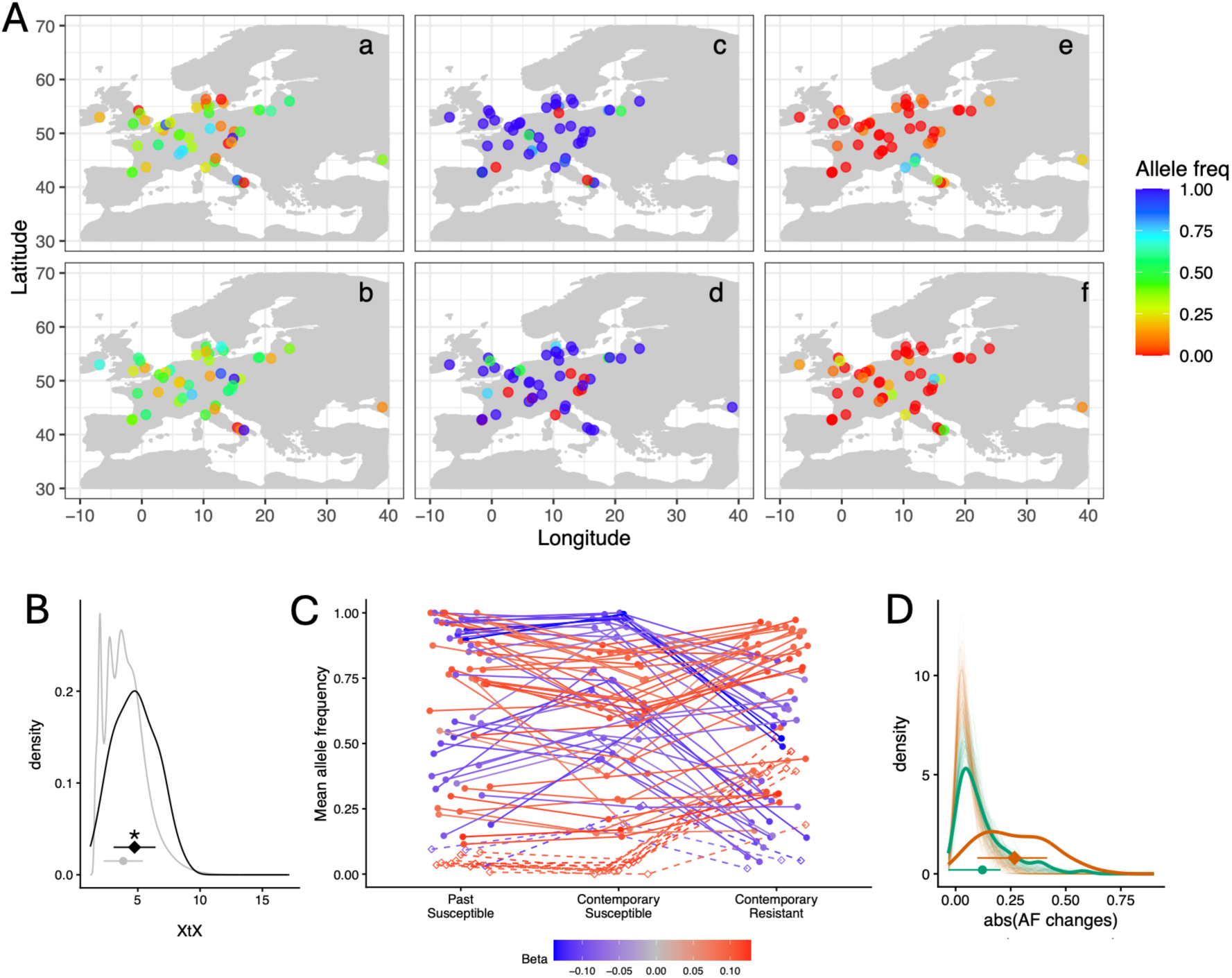
Recent NTSR polygenic evolution via large allele frequency changes in Europe. A. Geographical distribution of some of the loci identified as in association with herbicide resistance. Each panel refers to one SNP. Each dot represents one population, with colours representing the allele frequency of the alternative allele. B. XtX comparison between NTSR-associated loci (black) and the rest of the genome (grey) in historical susceptible populations. C. Alternative allele frequencies across time and resistance profile. Each dot represents one SNP. Colours are based on effect sizes (*Beta*) estimated per SNP in BayPass. Diamond dots connected by dashed lines represent SNPs with an average frequency of less than 0.1 in historical susceptible populations. D. Distribution of allele frequency changes (absolute values, AF). Orange marks changes between contemporary susceptible and resistant populations, while green represents changes in allele frequencies between susceptible historical and contemporary populations. The thicker lines represent the observed distribution for the 76 candidate loci identified, while the thinner lines represent the distribution of changes across 100 replicates of 76 randomly draw variants across the genome. In B. and D., the dots show median and the whiskers standard deviation, and the asterisks show statistically significant differences between the two distributions for the one-side Kolmogorov-Smirnov test.

Moreover, we found that most NTSR loci (83%) were already segregating at frequencies ≥ 0.1 in historical susceptible populations (Supplementary Figure 7), consistent with a role for standing genetic variation. Overall, our results indicate that NTSR has evolved recently in Europe, with a polygenic architecture, involving large effect variants. This evolutionary response to human imposed pressures was mainly driven by standing genetic variation present in historical populations, revealing a degree of pre-adaptation.

**Supplementary Figure 3.**
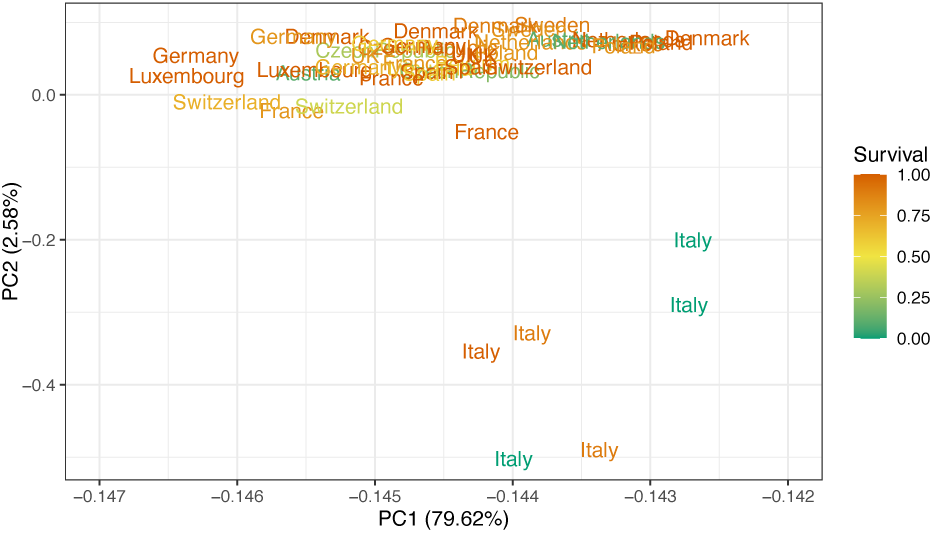
Whole-genome PCA for the 48 contemporary European populations. Each population is represented by its country of origin, coloured by survival, as shown in the legend. Computed in BayPass. Variance explained by each principal component is shown in parenthesis.

**Supplementary Figure 4.**
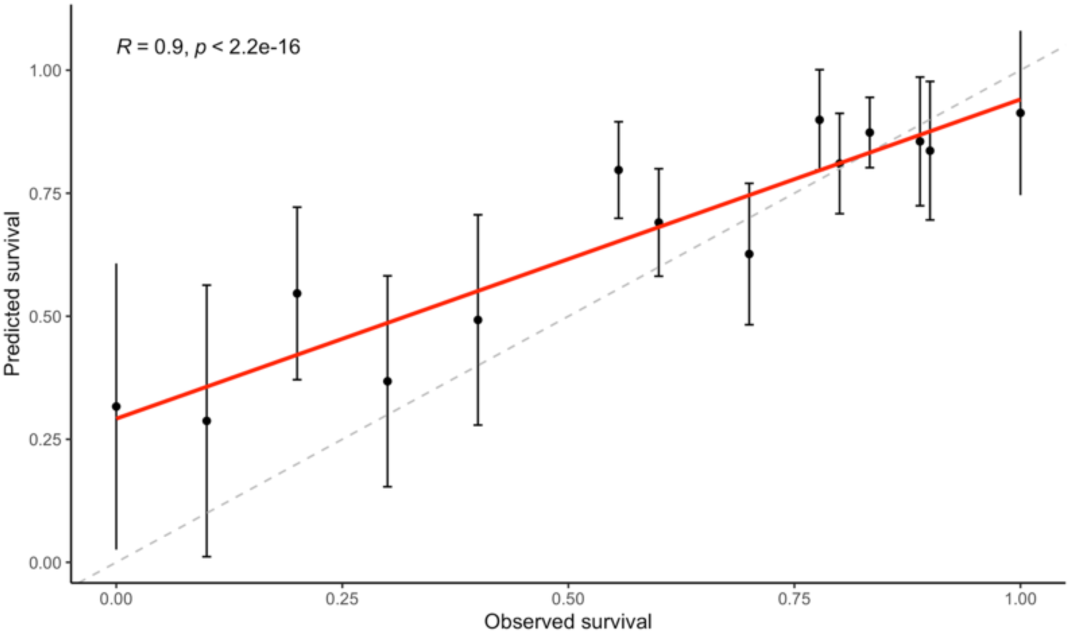
NTSR predictive model. Correlation plot between observed survival proportions (x-axis) on the 48 contemporary European blackgrass populations and the corresponding predicted values (y-axis). Results from a Bayesian generalized linear model based on the allele frequencies at the 76 candidate variants. Each dot represents the median across 100 iterations for the observed survival proportion, while the whiskers refer to the standard deviation. The grey dashed line marks the identity line, and the red the linear relationship between our predicted values. Correlation coefficient and p-value refer to a Spearman correlation.

**Supplementary Figure 5.**
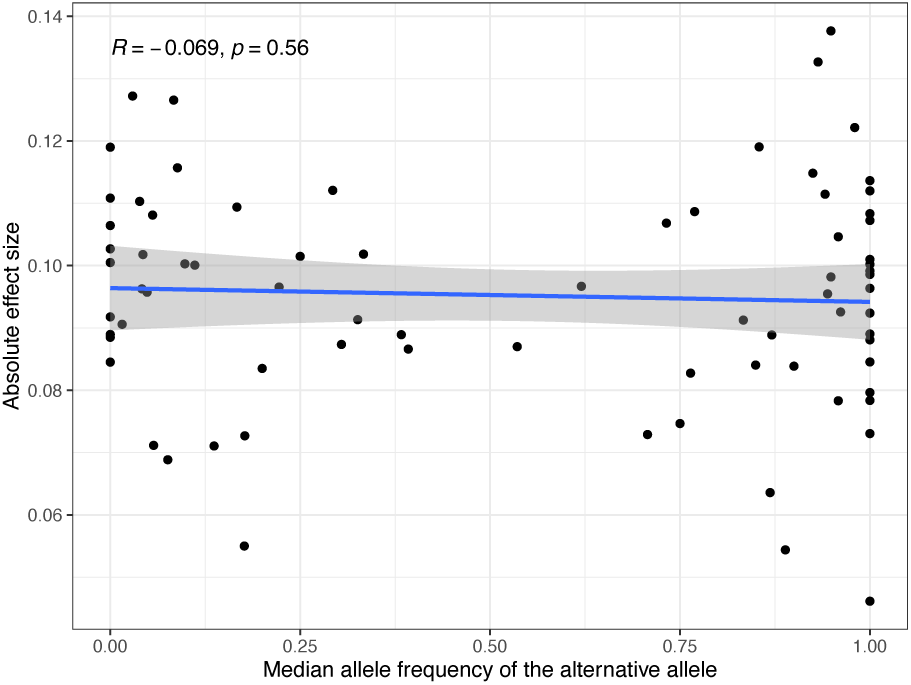
Correlation between median allele frequency across the 48 European populations (x-axis) and the absolute effect size (y-axis) of each NTSR-associated loci. Correlation coefficient and p-value from Spearman correlation.

**Supplementary Figure 6.**
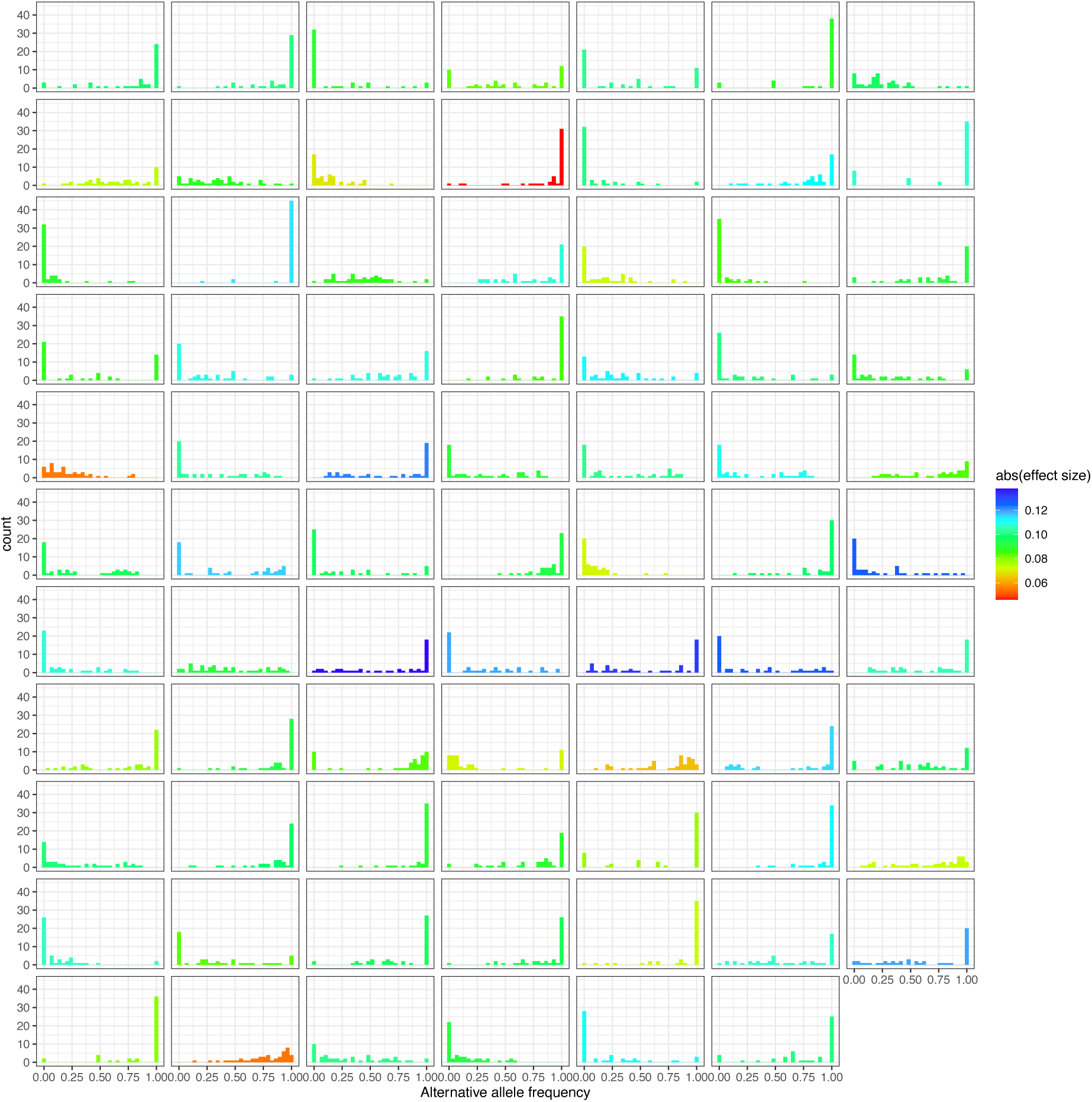
Allele frequency distribution of each NTSR-associated allele in the 48 European populations. Each panel refers to one representative SNP. X-axis shows the allele frequency of the alternative allele and y-axis the number of populations with the allele at that frequency. Colours represent the estimated effect size (in absolute values) for each associated SNP.

**Supplementary Figure 7.**
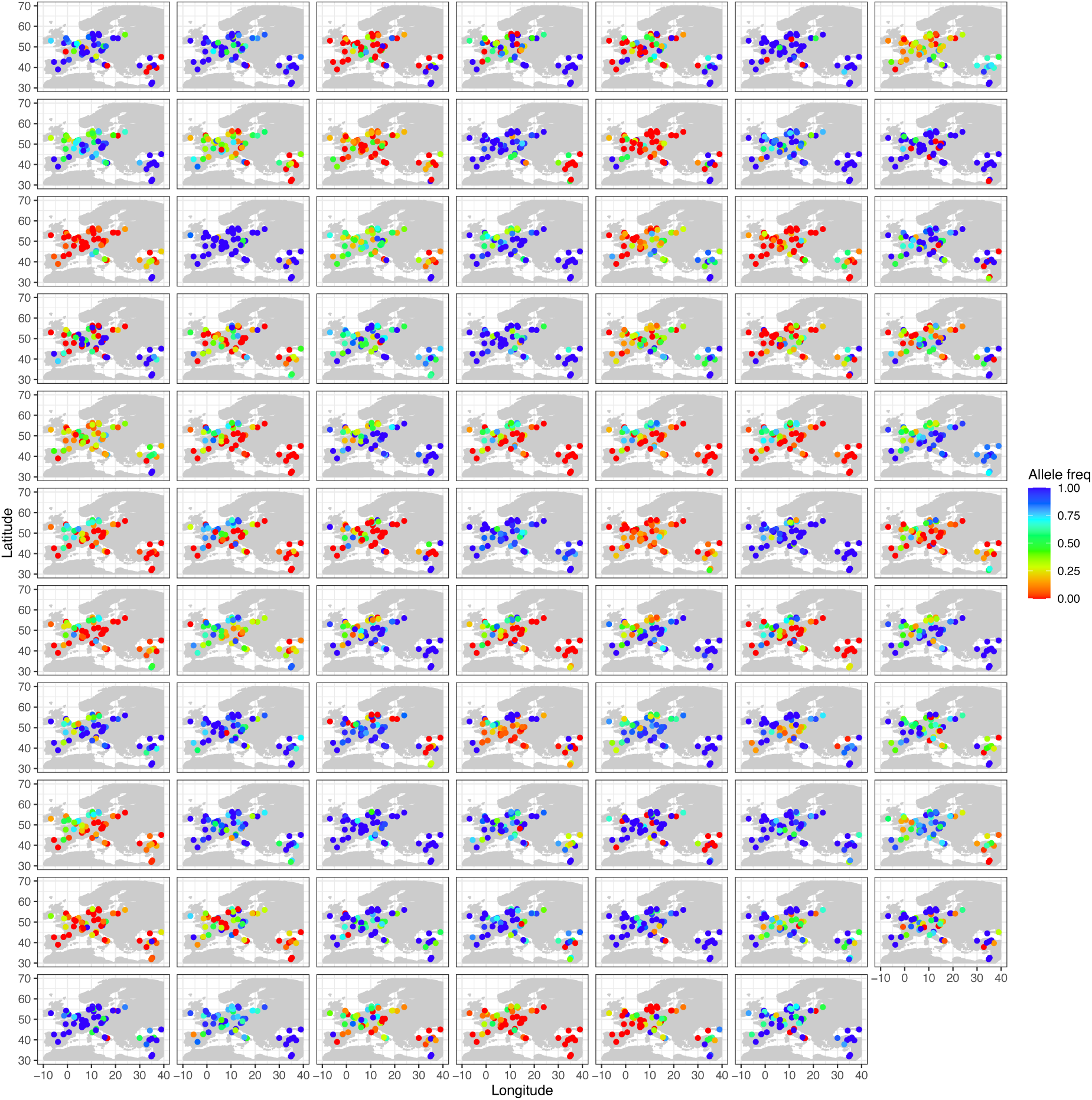
Geographical distribution of allele frequencies in the 64 populations sequenced. Each panel refers to one NTSR-associated SNP. Each dot represents one population, in their site of collection (x-axis shows longitude and y-axis latitude), coloured by alternative allele frequency of the SNP in question.

### NTSR evolved under strong directional selection

Under the hypothesis of pre-adaptation, we calculated XtX – a measure of adaptive differentiation – in resistance-associated loci in Middle Eastern historical populations, i.e., populations naïve to herbicide selection. NTSR-associated loci showed greater XtX values than genome-wide expectations (one-sided Kolmogorov–Smirnov test, D = 0.253, p = 0.01; Figure 3B), suggesting these loci may have been under selection before herbicide exposure.

In European contemporary populations, XtX values at NTSR-associated loci were also significantly higher than across the genome (one-sided Kolmogorov-Smirnov test, D = 0.499, p-value = 6.21×10^-6^; Supplementary Figure 8A), indicating these loci are also currently under positive selection. Moreover, candidate loci did not show the reduced diversity characteristic of hard sweeps (Watterson’s θ, one-sided Kolmogorov-Smirnov test, D = 0.007, p = 0.92; Supplementary Figure 8C), indicating that selection drove allele frequency shifts across multiple genetic backgrounds rather than fixation of a single allele, consistent with soft sweeps on standing genetic variation.

Finally, we reconstructed the evolutionary trajectory of NTSR in blackgrass by tracking allele frequency changes at the 76 resistance-associated loci. Assuming historical populations were susceptible – consistent with our data and the hypothesis that resistance evolved under recent human selection – we compared allele frequencies across historical susceptible, contemporary susceptible and contemporary resistant populations (Figure 3C and 3D, Supplementary Figure 9).

Frequency shifts between historical and contemporary susceptible populations were lower (mean |ΔAF|= 0.12) than between contemporary susceptible and resistant populations (mean |ΔAF|= 0.27, one-sided exact two-sample Kolmogorov-Smirnov test, D = 0.474, p-value < 2.2×10^-16^; Figure 3C and 3D), indicating that larger allele frequency changes occurred during recent evolution. A robustness analysis using 100 sets of 76 randomly selected variants with initial allele frequencies matching the candidate variants showed that the large allele frequency shifts observed far exceeded the null distribution expected under drift (two-sided exact two-sample Kolmogorov-Smirnov test, D = 0.637, p-value < 2.2×10^-16^; Figure 3D), indicating that these changes are unlikely to result from drift alone and are consistent with selection driving resistance.

The same pattern held for alleles that were rare in historical populations (AF < 0.1). Rare alleles showed larger allele frequency shifts between contemporary susceptible and resistant populations (mean |ΔAF| = 0.33,) than between susceptible historical and contemporary populations (mean |ΔAF| = 0.05; one-sided exact two-sample Kolmogorov-Smirnov test, D = 0.923, p-value < 2.2×10^-16^). Furthermore, rare alleles also showed higher estimated effect sizes in contemporary populations than more frequent alleles (one-sided two-sample Kolmogorov-Smirnov test, D = 0.275, p-value < 2.2×10^-16^; Figure 3C). These results suggest that both standing and historically rare variation contributed to the recent evolution of NTSR.

**Supplementary Figure 8.**
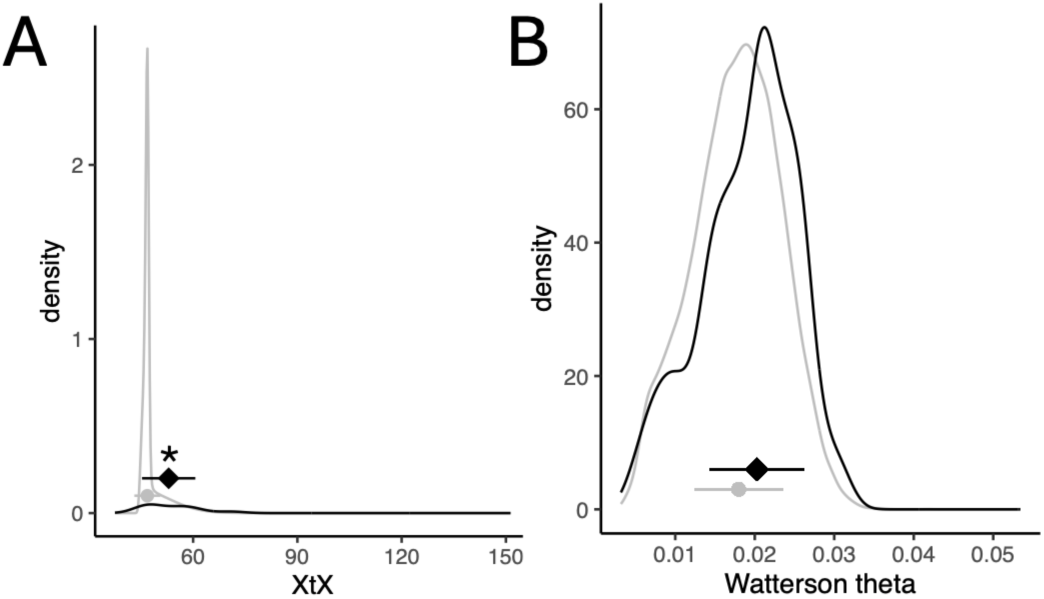
Positive selection in associated loci but no signal for hard sweeps in contemporary European populations. NTSR-associated loci (black) compared to the rest of the genome (grey) for A. XtX, and B. Watterson’s θ. Dots show the median and whiskers the standard deviation. Asterisks mark statistically significant differences between the two distributions, according to one-side Kolmogorov-Smirnov tests.

**Supplementary Figure 9.**
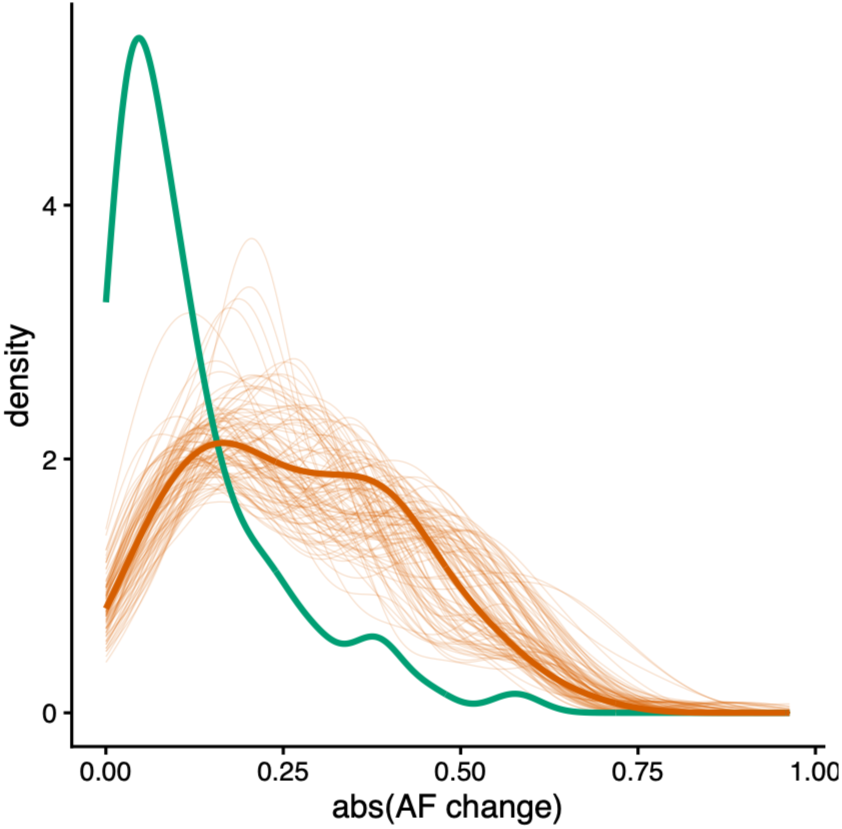
Allele frequency changes for the 76 candidate loci. A robustness analysis was done to assess the effect of different numbers of sampled populations. A downsample of contemporary resistant population was conducted by randomly selecting eight populations, 100 times. Allele frequency changes were calculated across categories as in Figure 3D.

### Gene amplification of glutathione S-transferases may underlie NTSR

The 76 QTLs we identified in GWAS (Figure 4A) overlapped with 30 annotated genes (Supplementary Table 2), including candidates with plausible roles in herbicide resistance, such as those involved in stress responses (ALOMY7G38755, ALOMY3G13663, ALOMY1G09524), detoxification (ALOMY7G38035, ALOMY1G09524), and pathogen defence (ALOMY7G38754). Notably, this set included three glutathione S-transferases (GSTs) (ALOMY3G13665, ALOMY3G13666, ALOMY3G13667), one UDP-glucosyltransferase (UGT) (ALOMY1G09524) and one very-long-chain 3-ketoacyl-CoA synthase (ALOMY1G09510), all key enzyme families previously implicated in NTSR (Cummins et al., 1999; Yuan et al., 2007; Délye, 2013; Cummins et al., 2013; Yu and Powles, 2014; Gaines et al., 2014, 2020).

The three GSTs identified here are clustered within a 7 kb region of the blackgrass genome (chr3: 237178000 – 237185000), with two additional GSTs (ALOMY3G13668 and ALOMY3G13670) located within 40 kb (Supplementary Figure 10A). These five GSTs share over 75% sequence identity (Supplementary Figure 10B and 10C). Notably, ALOMY3G13667 and ALOMY3G13670 were previously identified via RNA-seq as differently expressed genes linked to resistance in two strongly NTSR F2 recombinant blackgrass populations (Cai et al., 2023). Moreover, ALOMY3G13667, together with ALOMY3G13668 and ALOMY3G13670, has been implicated in detoxification of fenoxaprop and flufenacet, an herbicide inhibitor of the synthesis of very-long-chain fatty acids (VLCFAs) (Parcharidou et al., 2023). Interestingly, all five GSTs also co-localize with GSTU2, an *a priori* NTSR candidate (Cai et al., 2023).

Even after correcting for linkage disequilibrium, nine distinct QTLs were detected within this region in our GWAS, prompting a closer examination (Figure 4B). Sequence coverage analysis revealed pronounced differences between resistant and susceptible populations across the region. Normalized per-pool coverage was significantly correlated with herbicide survival across the eleven 1-kb windows examined (chr3:237175000-237185000) (Figure 4B, Supplementary Figure 11). In resistant populations, normalized coverage increased up to 10-fold (window 237184000), and coincided with QTL peaks, suggesting the presence of structural variation, likely copy-number variation, associated with resistance. Notably, these coverage peaks align with the three candidate GSTs, suggesting that gene amplification of these loci may underlie NTSR in blackgrass, consistent with previously reported expression differences between resistant and susceptible individuals at these loci (Cai et al., 2023; Parcharidou et al., 2023).

Interestingly, these extreme fold-changes occurred exclusively in contemporary European populations, were more localized in Scandinavia and western Europe, and were relatively rare (only 5 populations out of the 64 sequenced showed an increase above 10-fold; Figure 4C), suggesting this could be a recent amplification event.

**Figure 4.**
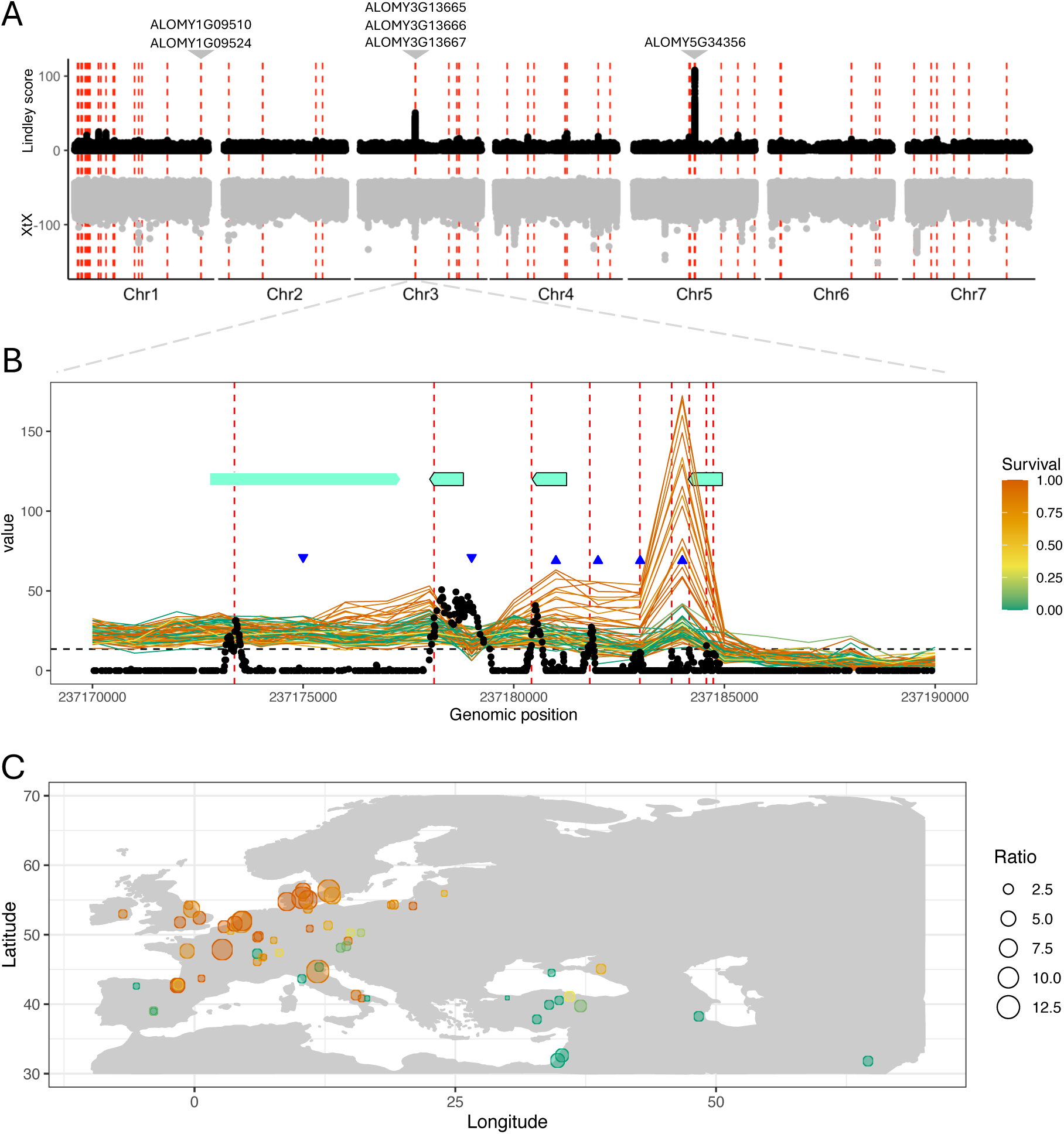
GSTs as candidate genes to underlie NTSR in blackgrass. A. Miami plot showing genome-wide association with survival in the subset of 48 European populations (top, in black; y-axis shows Lindley values from the local score approach) and XtX values (bottom, in grey; negative values used just for plotting), across the genome (x-axis). Red vertical dashed lines represent significant regions identified with the local score approach. On top, some candidate genes are highlighted, including the three GSTs identified. B. Zoom in on the region in association in chromosome 3 underlying nine independent significant regions. X-axis represents genomic position and y-axis value for different metrics: black dots replicate the Lindley score from the Manhattan plot above; colourful lines correspond to per-population coverage in 1-kb windows. The colour matches the survival (herbicide resistance) of the corresponding population. Vertical red lines highlight the significantly associated regions. Horizontal dashed line marks mean coverage across chromosome 3, across all populations. The horizontal light green bars represent genes annotated in this region, with the three GSTs highlighted by the black lines around the boxes (same GSTs highlighted in A). The blue arrows mark windows with normalized coverage significantly correlated with survival (up: positive correlation, down: negative correlation, data from Supplementary Figure 11). C. Geographical distribution of normalized increased coverage in window chr3:237184000. Each dot marks a population, coloured by resistance (survival). The size of the dot (*Ratio* in the legend) is proportional to the fold change in this window compared to the mean coverage of chromosome 3 for each population.

**Supplementary Figure 10.**
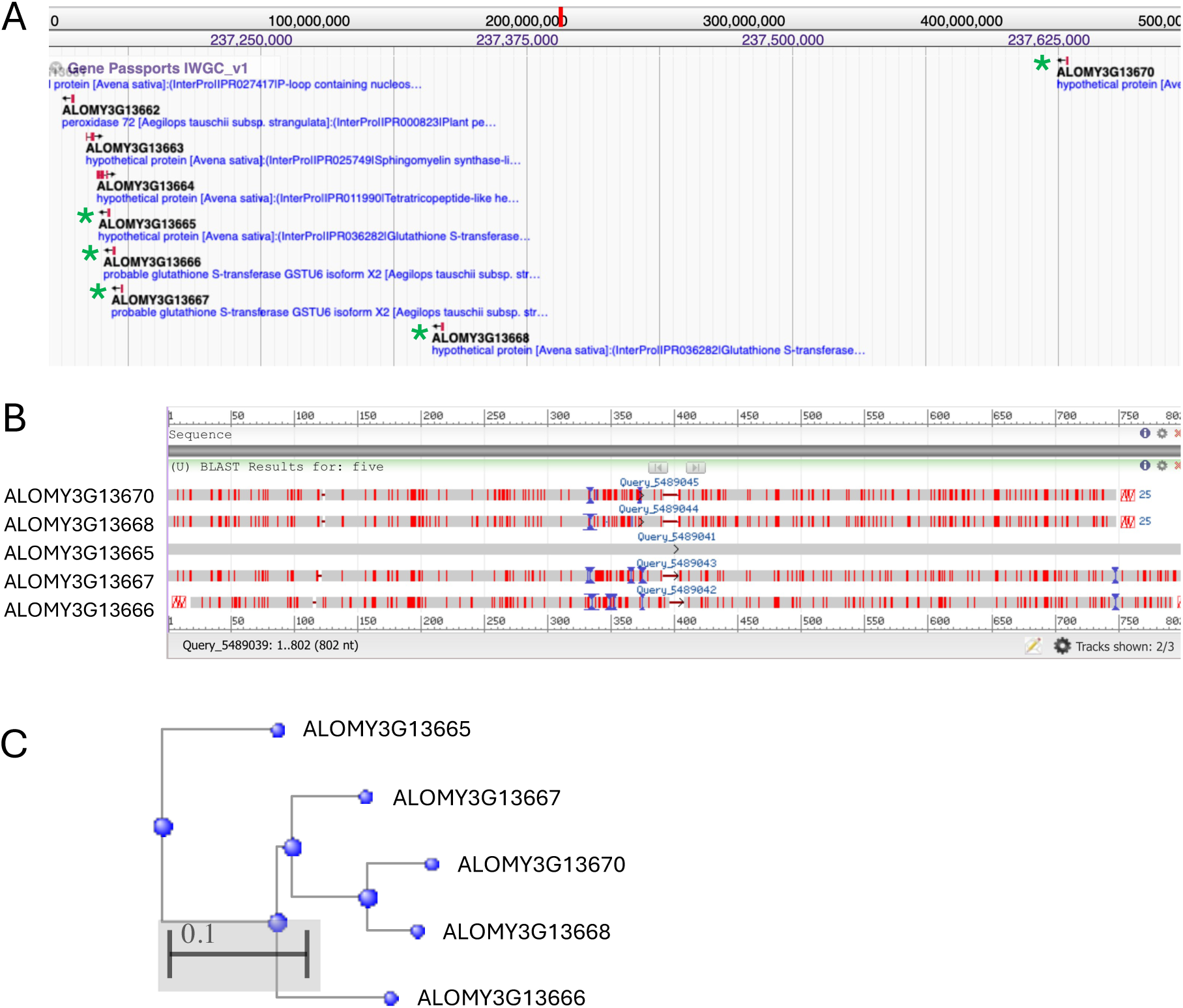
Zoom-in on five co-localizing GSTs in the blackgrass genome. A. Zoom-in on the blackgrass genome at region chr3:237130000-237690000 from the GenomeBrowser in the IWGC WeedPedia (https://weedpedia.weedgenomics.org). The five GSTs are highlighted by the green asterisks. B. Alignment and sequence similarity between the five GSTs. Results from BLAST in NCBI (https://blast.ncbi.nlm.nih.gov/Blast.cgi). C. Similarity tree computed in BLAST, using the Fast Minimum Evolution method.

**Supplementary Figure 11.**
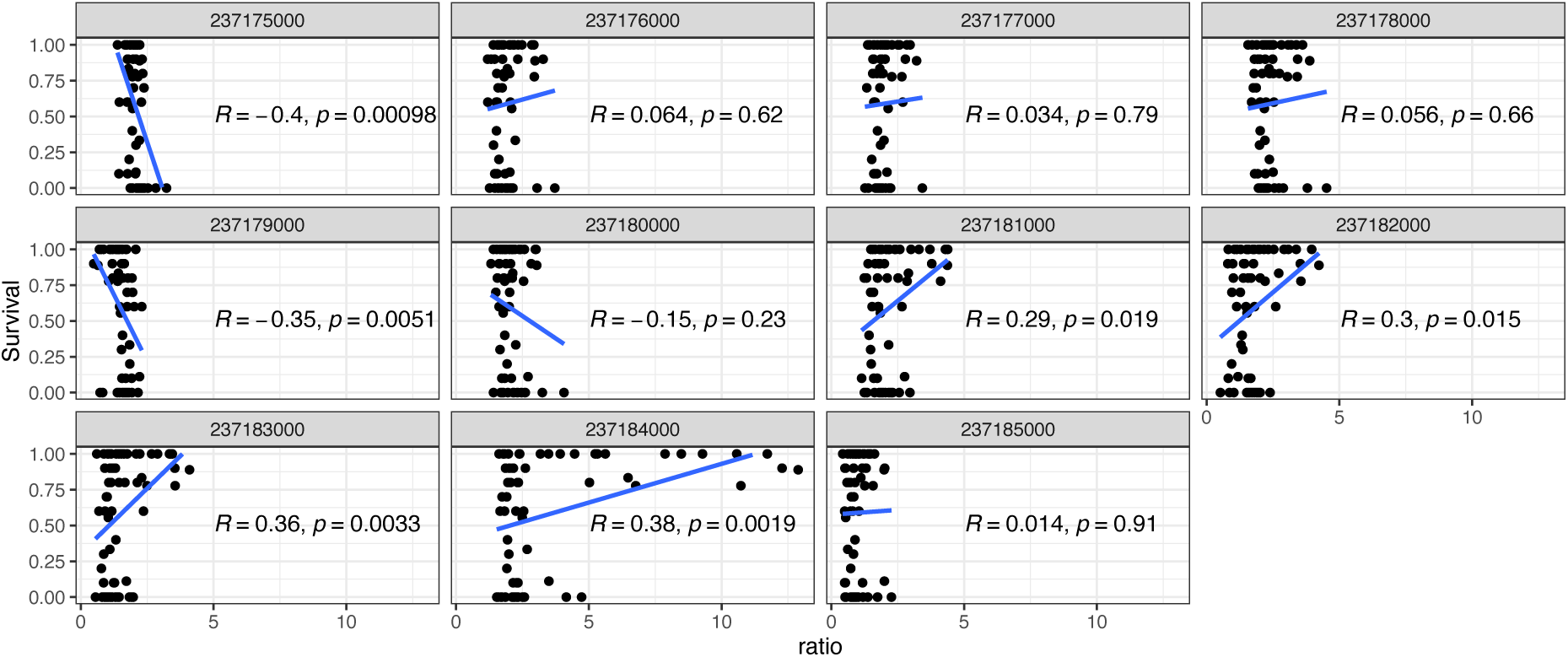
Pearson correlation between fold change in coverage (x-axis) and survival (y-axis). Each panel corresponds to one of the eleven 1-kb windows considered in the region of interest in chromosome 3, labelled at the top. Each dot corresponds to one population. *Ratio* (x-axis) refers to the fold change in coverage for each 1-kb window relative to the mean coverage of the corresponding population across chromosome 3.

## Discussion

Combining phenotypic assays with whole-genome sequencing of diverse blackgrass populations, we show that herbicide resistance evolved recently and rapidly through monogenic and polygenic architectures on standing genetic variation. Crucially, inclusion of populations collected prior to widespread herbicide use allowed us to directly trace the resistance alleles evolutionary trajectory from pre-selection baselines. Moreover, this study represents the first genome-wide analysis of the genetic basis of herbicide resistance in natural blackgrass populations and one of the few conducted in major herbicide resistant weeds. Our analysis offers comprehensive insights into the genomic architecture of contemporary adaptation under recent, intense anthropogenic selection.

### Large-effect monogenic vs polygenic architectures

Herbicide resistance reflects a combination of monogenic TSR and polygenic NTSR. Consistent with previous studies (Comont et al., 2020; Kersten et al., 2023), we show ACCase resistance is conferred by co-occurring TSR and NTSR mechanisms.

Eight known, previously characterized TSR mutations explained 45% of the phenotypic variance in resistance, while polygenic NTSR was supported by 76 genomic regions associated with resistance. Our GWAS in natural populations identified substantially more NTSR-associated loci than identified in recombinant mapping populations (Cai et al., 2023). Similar patterns have been reported in *Amaranthus tuberculatus* (Kreiner et al., 2021) and *Ipomoea purpurea* (Van Etten et al., 2020), where >250 and ∼80 loci, respectively, contribute to metabolic resistance, indicating that highly polygenic architectures for NTSR are common across weed species.

While TSR mutations confer robust resistance, consistent with classical hard sweep dynamics (Powles and Yu, 2010; Pritchard et al., 2010; Gaines et al., 2020), NTSR is characterized by a polygenic architecture, involving numerous loci of moderate-to-large-effect (Barghi et al., 2020; Boyle et al., 2017; Hermisson and Pennings, 2005). These loci include both widespread and population-specific variants. In *Ipomoea purpurea*, some resistance-associated loci showed parallel signatures of selection among highly resistant populations, whereas others diverged among populations (Van Etten et al., 2020). Such polygenic and partially population-specific architectures allow incremental adaptation to heterogeneous selective pressures. This is reflected in the geographical distribution of NTSR alleles: some variants are shared among populations, whereas many are unique, producing a continuous, cumulative, “dimmer-like” response rather than the discrete “switch-like” effects typical of TSR. This pattern is consistent with models of adaptation from standing genetic variation, in which resistance evolves through simultaneous selection on many loci rather than reliance on a single mutation (Barghi et al., 2020; Délye, 2013; Gaines et al., 2020; Höllinger et al., 2019; Pritchard et al., 2010).

### Standing genetic variation vs *de novo* mutations and the rate of evolution

We detected no *ACCase* TSR mutations in historical populations, suggesting these were absent or very rare (below our detection threshold) prior to herbicide use, and subsequently increased in frequency under recent, strong herbicide selection.

However, as our pool-seq design included only 25 individuals per population, our power to detect rare alleles is limited and we cannot exclude the possibility that TSR variants were present at very low frequencies as standing variation. Previous work supports this possibility: in *Amaranthus tuberculatus*, age estimates of TSR mutations predated herbicide use but remained rare until herbicide selection (Kreiner et al., 2022b), while in blackgrass, simulations showed rare standing TSR alleles could reach moderate frequencies within 30 generations under strong selection (Kersten et al., 2023).

Moreover, herbarium screenings found an *ACCase* Ile1781Leu mutation in blackgrass collected in 1888 (Délye et al., 2013a).

In contrast, most NTSR loci appear to predate herbicide use, segregating at moderate frequencies in historical susceptible populations, i.e., populations naïve to herbicide selection. These results indicate that herbicide resistance evolved from standing genetic variation – an inference only possible by the inclusion of live, pre-selection populations. Historical germplasm – especially when direct phenotypic validation is possible – does however carry inherent limitations, including the reduced number of populations available and undefined sampling protocols. Nonetheless, when these are accounted for, it represents an irreplaceable resource for reconstructing the evolutionary history of resistance at the genetic and phenotypic level, going beyond what herbarium specimens alone can offer (Délye et al., 2013a; Kreiner et al., 2022a).

Resistance-associated loci showed signatures of positive selection in historical, pre-herbicide populations, suggesting herbicides did not create resistance so much as recruit pre-existing adaptive variation. These loci were maintained by selection unrelated to herbicide stress, possibly through pleiotropic effects on other adaptations (Hawkins et al., 2019), supporting a “pre-adaptation” hypothesis, whereby resistance-prone weeds harbour standing variation in stress and defence pathways shaped by historical environmental pressures (Gaines et al., 2020; Powles and Yu, 2010).

Consistent with this idea, transcriptomic studies in blackgrass show overlap between metabolic NTSR and waterlogging stress responses (Harrison et al., 2024), and in *Amaranthus tuberculatus*, historical selection on *PPO* predates herbicide use and coincides with long-term agricultural adaptation (Kreiner et al., 2022a). Comparative genomics further suggests that gene families associated with detoxification and abiotic stress are unusually diverse in weeds. The blackgrass genome shows expansion of these gene families (Cai et al., 2023), while in *Leptochloa chinensis* and *Echinochloa crus-galli* polyploidization retained genes linked to abiotic stress and detoxification but lost disease resistance gene families (Ye et al., 2020; Chen et al., 2023). Together, these patterns suggest that weed evolution favours maintenance and diversification of stress and detoxification pathways, generating functional redundancy and standing variation that can later facilitate rapid adaptation to new selection pressures such as from herbicides (Lynch and Conery, 2000).

These findings align with theoretical models predicting rapid adaptation from standing variation, especially for polygenic traits. Strong directional selection acting on many loci can produce substantial phenotypic change within a few generations (Barrett and Schluter, 2008; Walsh and Lynch, 2018), shifting allele frequencies across numerous loci rather than relying on fixation of a single mutation (Hermisson and Pennings, 2005; Messer and Petrov, 2013; Höllinger et al., 2019).

### Repeatability and predictability

Geographical patterns in *ACCase* mutations likely reflect a combination of stochastic processes, variation in selection pressures, and fitness costs. Mutations conferring broad cross-resistance to ACCase herbicides have been repeatedly selected (Délye et al., 2008; Kaundun et al., 2013) and generally show little or no fitness penalty (Vila-Aiub et al., 2005; Menchari et al., 2008; Délye et al., 2013b; Vila-Aiub et al., 2015). In contrast, less frequent mutations often impose stronger fitness costs, that limit their spread (Menchari et al., 2008; Vila-Aiub et al., 2015; Du et al., 2017).

Most *ACCase* mutations occurred in diverse genomic backgrounds, suggesting repeated and parallel origins. However, some showed evidence of a single origin followed by dispersal, and others a combination of both processes. Similar patterns have been reported in blackgrass, and other grasses, where identical amino-acid substitution arise independently under herbicide selection (Powles and Yu, 2010; Wang et al., 2010; Alarcón-Reverte et al., 2013; Baucom, 2016; Du et al., 2017, 2017; Gaines et al., 2020; Vázquez-García et al., 2020; Kreiner et al., 2022b; Kersten et al., 2023).

In contrast, NTSR showed limited repeatability at individual loci despite strong phenotypic predictability. The 76 NTSR-associated SNPs predicted resistance effectively across populations, indicating they capture a substantial fraction of phenotypic variation. However, most loci varied widely in frequency among populations, a pattern also observed in *I. purpurea* (Van Etten et al., 2020). These combinations of strong phenotypic predictability but limited genetic sharing are characteristic of polygenic adaptation, where similar phenotypes arise through different allele combinations at many loci. This contrasts with TSR, which often shows classic parallel evolution with repeated amino-acid substitutions under strong selection (Baucom, 2016). Together, these results illustrate how the predictability of adaptation depends on genetic architecture and highlight the need to consider both monogenic and polygenic components when forecasting evolutionary responses.

### Genetic basis of NTSR

Consistent with previous work (Délye, 2013; Gaines et al., 2020), NTSR appears to evolve through modifications in broad stress-response pathways. Resistance-associated loci included glutathione S-transferases (GSTs), UDP-glucosyltransferases (UGTs), and stress signalling genes families repeatedly implicated in NTSR across species in genetic mapping, transcriptomic and functional studies (Shimabukuro et al., 1970; Anderson and Gronwald, 1991; Van Etten et al., 2020; Tal et al., 1993; Goggin et al., 2021; Cai et al., 2023; Parcharidou et al., 2023; Gupta et al., 2023).

Three co-localized GSTs underlying a major QTL represent strong functional NTSR candidates. Increased read coverage in resistant populations suggests possible copy number variation. Amplification of detoxification genes is a well-documented mechanism of herbicide resistance, as additional gene copies elevate transcript abundance and enzymatic activity, enhancing metabolic detoxification and reducing herbicide accumulation (Gaines et al., 2020). Amplification of *EPSPS* (5-enolpyruvylshikimate-3-phosphate synthase) in glyphosate-resistant weeds provides a classic example, with copy numbers ranging from 3 to 160 across at least 12 species (Wiersma et al., 2015; Salas et al., 2015; Patterson et al., 2019; Gaines et al., 2020).

Our results show that herbicide resistance in *Alopecurus myosuroides* represents a striking case of rapid contemporary adaptation driven by ∼50 years of intense anthropogenic selection. By integrating phenotypic assays with whole-genome sequencing across diverse germplasm, we demonstrate that resistance arises through a combination of monogenic and polygenic mechanisms, dominated by polygenic adaptation via soft sweeps on standing genetic variation. Defence– and stress-related pathways provide reservoirs of pre-adaptive variation that can be repeatedly recruited under novel chemical stress. Consequently, resistance has evolved independently across regions through different allelic combinations, producing strong phenotypic repeatability despite genetic heterogeneity. By reconstructing evolutionary trajectories across space and time, this study addresses long-standing questions about the architecture, sources of variation, and predictability of rapid adaptation, establishing blackgrass resistance as a powerful model of evolution under strong anthropogenic selection.

## Acknowledgments

The authors thank colleagues who collected and shared blackgrass seeds, namely Alberto Collavo, Bayer Crop Sciences, Germany; David Comont, Rothamsted Research, UK; Husrev Mennan, Ondokuz Mayis University, Turkey; Katěrina Hamouzová, Czech University of Life Sciences, Czech Republic; Donato Loddo, Institute for Sustainable Plant Protection, National Research Council, Italy; Joel Torra Farré, Universitat de Lleida, Spain; Mette Sønderskov, Aarhus University, Denmark; Maor Matzrafi, Volcani Institute, Israel; and the United States Department of Agriculture Germplasm Resources Information Unit (GRIN), USA. We are also grateful to glasshouse technical staff at the Taastrup campus of the University of Copenhagen for help with maintenance of plants. The project was supported by the Novo Nordisk Foundation (Grant number: NNF21OC0068600). The funders had no role in study design, data collection and analysis, decision to publish, or preparation of the manuscript.

## Competing interests

The authors declare no conflict of interests.

## Author contributions

CN designed the study, conducted all analyses, wrote the first draft of the manuscript, and edited subsequent drafts. AB collected phenotypic data. PN obtained seed material, designed the study, and contributed to writing and editing of manuscript drafts.

## Data availability

Genomic data are available on NCBI SRA under Bioproject PRJNA1285609. All code used in bioinformatic processing, analyses and data visualization is available in the GitHub repository https://github.com/celianeto/blackgrass_HR.

## Notes

### Competing Interest Statement

The authors have declared no competing interest.

